# NVUAtlas: A Comprehensive Single-Nucleus RNA-Seq Resource for the Human Neurovascular Unit in Alzheimer’s Disease

**DOI:** 10.64898/2025.12.03.692162

**Authors:** Xiaojia Tang, Douglas A Nelson, Mikel Hernaez, Karunya K Kandimalla, Krishna R Kalari

**Affiliations:** Department of Quantitative Health Sciences, Mayo Clinic, Rochester, MN, USA; Department of Pharmaceutics and Brain Barriers Research Center, University of 6 Minnesota, College of Pharmacy, Minneapolis, MN, USA; Centro de Investigación Médica Aplicada (CIMA), Universidad de Navarra, Spain

**Author notes:** **Corresponding author: Krishna R Kalari**, Department of Quantitative Health Sciences, Mayo Clinic, Rochester, MN, USA.

## Abstract

**Background:** Neurovascular unit (NVU) dysfunction is being recognized as one of the earliest contributors to Alzheimer’s disease (AD) pathogenesis. However, systematic investigation of NVU dysfunction is currently limited by lack of access to molecular level information due to underrepresentation of vascular and mural cells in standard single-nucleus RNA sequencing (snRNA-seq) datasets. Consequently, existing transcriptomic atlases lack the resolution necessary to capture the coordinated intercellular signaling and dysfunction across vascular components of NVU, including endothelial cells and pericytes.

**Methods:** We constructed the Human NVU Atlas by integrating 11 publicly available snRNA-seq datasets, including vascular-enriched samples, from the human prefrontal cortex. This comprehensive dataset aggregates over 4.2 million nuclei from 748 donors, including AD patients and age-matched controls. We utilized a unified probabilistic pipeline based on deep generative models (SCVI) to perform batch-aware integration and employed an ensemble of supervised and deep-learning classifiers to rigorously re-annotate cell types. Differential expression and ligand-receptor interaction analyses were subsequently performed to identify cell-type-specific disruptions in males versus females.

**Results:** The atlas successfully curated vascular populations from 11 studies to assemble the largest publically available NVU cohort of endothelial cells (2.8%) and pericytes (1.9%) alongside astrocytes and neurons. Differential expression analysis revealed that while neurons predominantly exhibited gene downregulation in AD, vascular cells displayed a pattern of transcriptional hyperactivity with significant gene upregulation. We also identified pronounced sex-specific vulnerabilities; females exhibited distinct inflammatory signatures and downregulation of basement membrane collagen genes (e.g., *COL4A1*, *COL4A2*) in pericytes, whereas these changes were not observed in males. Moreover, cell-cell interaction analysis revealed a widespread loss of collagen-integrin signaling between pericytes and neurons, suggesting the involvement of extracellular matrix disruptions in NVU dysfunction observed in AD.

**Conclusion:** The Human NVU Atlas provides a high-resolution, integrated transcriptomic framework for dissecting the cellular heterogeneity of the neurovascular unit. By uncovering sex-specific vascular mechanisms and disrupted intercellular communication, this resource highlights the critical role of vascular cells in AD progression and serves as a foundational reference for investigating cerebrovascular contribution to AD.

## INTRODUCTION

The neurovascular unit (NVU) is the functionally integrated multicellular unit characterized in this atlas, consisting of endothelial cells, pericytes, astrocytes, neurons. Endothelial cells form the capillary wall and constitute the blood-brain barrier (BBB), which regulates nutrient delivery and waste clearance through tight junctions and polarized transport mechanisms. (1) Pericytes, which are particularly dense in the brain, wrap around the abluminal surface of capillary endothelium, where they regulate capillary diameter, support the specialized endothelial phenotype of the BBB, and modulate angiogenesis (2,3). Astrocyte endfeet closely surround the vasculature, regulating extracellular ion and water homeostasis and providing metabolic support to the NVU (4). Neuronal processes course along capillaries, establishing close anatomical proximity to NVU cells. This dense cellular architecture is brought together by the basement membrane, an extracellular matrix that envelopes pericytes and anchors astrocyte endfeet, providing not only structural cohesion but also functioning as a hub for NVU paracrine signaling and mechanotransduction(5).

The NVU is not merely a structural assembly but a dynamic interdependent system. On short timescales, the NVU mediates neurovascular coupling, in which neuronal activity triggers signaling cascades that engage astrocytes, pericytes, and endothelial cells to regulate blood flow and deliver metabolic support (6). Over longer timescales, NVU cells engage in interdependent metabolic, growth factor, and inflammatory signaling to maintain NVU homeostasis, preserve barrier integrity, and support neuronal function(7). Thus, dysfunction of one or more NVU components can lead to broad functional disruptions contributing to neurological disease (8).

Disruption of the NVU is recognized as an early and significant contributor to Alzheimer’s disease pathogenesis, with BBB dysfunction often preceding neurodegeneration and cognitive decline (9,10). Vascular pathology is a common comorbidity of AD, with studies reporting cerebrovascular lesions in two-thirds of autopsy-confirmed AD cases (11,12). Across the NVU, multiple cell types exhibit structural and functional impairments. Astrocytes display a reactive phenotype that impairs their metabolic support of the NVU, compromises BBB integrity, and promotes vascular inflammation (13). BBB endothelial cell dysfunction impairs a wide range of transport and signaling processes, including glucose uptake into the brain and clearance of amyloid-β peptides (14,15). Pericytes, the main supporting cells of the microvascular endothelium, exhibit degeneration and detachment in AD, which can lead to cerebral hypoperfusion and loss of BBB integrity (16,17).The basement membrane becomes fragmented, fibrotic, and compositionally altered in AD due to dysregulated expression and turnover of extracellular matrix proteins, disrupting NVU paracrine signaling and structural coherence (18–20).

The primary methods currently used for transcriptomic investigation of the human brain include bulk RNA sequencing, single-cell (scRNA-seq), single-nucleus RNA sequencing (snRNA-seq), and spatial transcriptomics (21). Bulk RNA-seq provides an aggregate transcriptome profile derived from all cell types present in a homogenized tissue sample, with contributions of each cell type weighted by their relative abundance (21). Single-cell and single-nucleus RNA-seq enable high-resolution profiling of individual cell types and states but rely on tissue dissociation or nuclear isolation, which can distort cellular proportions and introduce technical artifacts such as transcriptional dropout (22,23). In postmortem brain tissue, snRNA-seq is often the preferred approach due to its compatibility with frozen material and its avoidance of live-cell dissociation, which reduces transcriptional artifacts caused by mechanical stress. At the same time, snRNA-Seq does not capture cytoplasmic transcripts or local mRNA pools, a limitation more relevant to neurons than vascular cell types (21). Spatial transcriptomic methods preserve tissue architecture and enable spatial mapping of gene expression but are typically limited to multicellular resolution, requiring computational deconvolution, or are prohibitively expensive and low-throughput (21), limiting their practical application to NVU-focused studies across large AD cohorts. In single-nucleus datasets, neurons typically dominate, not only because of their relative abundance in brain tissue but also because standard nuclear isolation protocols tend to favor neuronal recovery and lead to loss of vascular nuclei (24). As a result, vascular and perivascular cells, particularly endothelial cells and pericytes, are disproportionately underrepresented, with sparse or inconsistent annotations and unreliable counts (25,26).

Current AD-focused large-scale transcriptomic studies, including ROSMAP(27), SEA-AD(28), ssREAD(29), and TACA(30), have advanced the field considerably, each contributing unique datasets and perspectives. For example, SEA-AD integrates multimodal data to map brain cell types and their alterations in Alzheimer’s disease, while ssREAD provides broad access to single-cell, single-nucleus, and spatial transcriptomic datasets across multiple brain regions and conditions (28,29). However, all these resources currently offer limited resolution of vascular and mural cells of the NVU. By contrast, the NVU Atlas integrates vascular-enriched and standard single-nucleus datasets from the human prefrontal cortex using a unified pipeline and annotation strategy for resolving all major NVU cell types. To our knowledge, no existing integration of AD transcriptomic datasets supports systematic investigation of the NVU at the necessary resolution and scale. This is due to both the underrepresentation of vascular populations in standard protocols and the absence of integration strategies designed explicitly for NVU-focused analysis across cohorts.

Beyond direct NVU analysis and hypothesis generation, we envision the NVU Atlas as a reference for additional applications, such as improved deconvolution of spatial transcriptomics and even bulk RNA-seq deconvolution. An interactive web-based interface is under development to support community access and exploration.

## METHODS

### Constructing the Human NVU Atlas for Alzheimer’s disease research

The Human NVU Atlas in our study is dedicated to the Alzheimer’s Disease research and specifically targets transcriptomic data from the pre-frontal cortex. We have compiled data from 11 publicly available cohorts, which are hosted on the GEO, Synapse, and CellxGene databases. The collected transcriptome data utilized droplet-based 10X 3’ sequencing technology, with majority of the datasets consisting of author-provided cell-level annotations. As summarized in Table 1, our Atlas consists of data from 748 patients, comprising 373 female and 375 male subjects, both AD patients and control individuals. Overall, we gathered approximately 4.25 million nuclei, with author-provided annotations available for 79.5% of these nuclei.

**Table 1:**
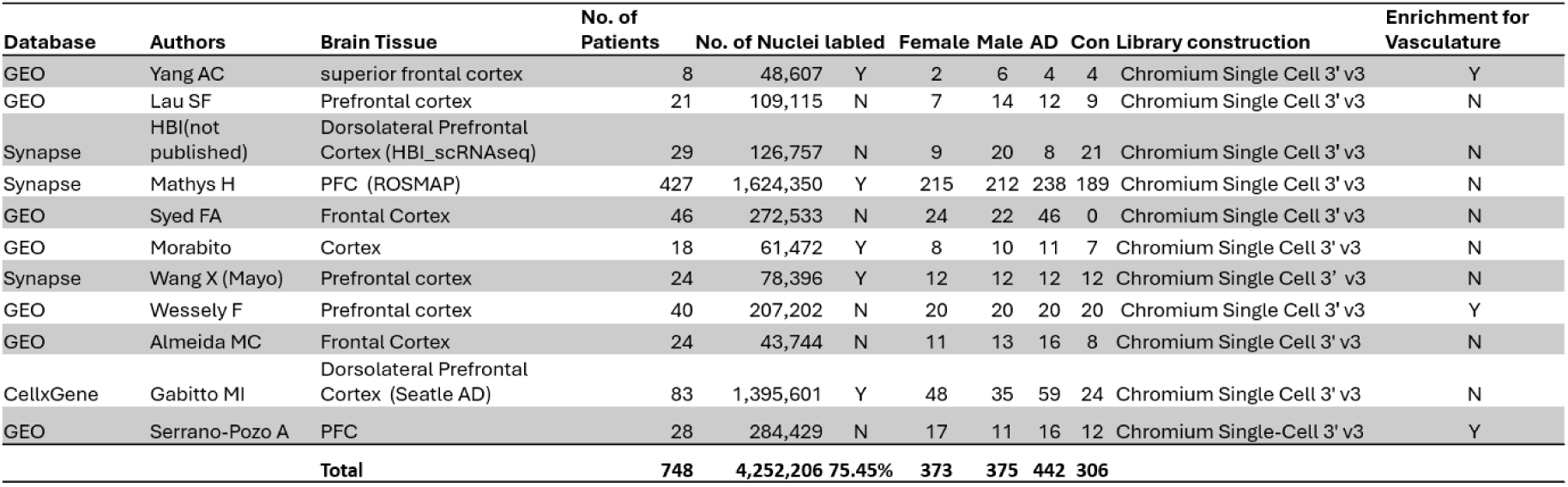
The table summarizes the extensive data compiled from multiple sources to construct the Human NVU Atlas, targeting Alzheimer’s Disease research with a focus on the frontal cortex. It shows the diversity and scale of the datasets integrated, detailing the number of patients, nuclei count, gender distribution, and the library construction techniques used.

### Optimizing gene expression data for integration using highly variable genes

To prepare the raw count data for integration, we first extracted raw counts from each of the studies and then processed each dataset independently. To address batch-specific variations in gene expression, which can impact integration performance, we utilized the highly_variable_genes() function with the “seurat_v3” option, as recommended by Single-cell Variational Inference (SCVI) tools (reference). This method helped us identify the top 2,000 highly variable genes (HVGs) across all datasets. We focused exclusively on these HVGs during the subsequent steps of atlas construction (integration) and cell type annotation.

### Integration and expansion of single-cell RNA-seq data using SCVI and scArches

We integrated the data from all cohorts using the SCVI tool, a state-of-the-art method for handling single-cell transcriptomics data. Initially, raw count matrices of the top 2000 HVGs from each study cohort were concatenated, along with study (batch) and other metadata. We then used the run_autotune() function from the SCVI package to identify optimal parameters for analyzing this concatenated dataset globally. The selected parameters for our model included a gene likelihood modeled with a Zero-Inflated Negative Binomial (ZINB) distribution, two hidden layers with 256 nodes each, and a latent space dimensionality of 40. The model underwent training for 200 epochs using these optimized parameters. Following training, we evaluated the latent representation and learned embeddings of each cell for the integrated data. This latent representation was then employed in downstream processes, such as cell type annotation analysis, while the embeddings were used for further visualization and analysis.

Furthermore, the integration framework supports the inclusion of new datasets into the existing integrated data cohort. Using the scArches method from the SCVI package, new single-cell RNA-seq datasets are treated as query sets with the main atlas cohort acting as the reference set. A model, pre-trained in the reference set, is used to load the query set, which is then trained with a weight decay of 0.0. This approach ensures that the latent representation of reference cells remains unchanged while incorporating new data through the query model.

### Cell-type reannotation and NVU Atlas development

We began by standardizing the author-provided cell type annotations from each study into major cell categories, including neurons, astrocytes, endothelial cells, pericytes, oligodendrocytes, OPCs, immune cells (specifically microglia), and categories for unknown cell types. Among all the input cells, 24.55% of cells were initially categorized as unknown. To address this, we employed the latent representation generated by the SCVI model to reannotate these unknown cell types. We utilized four different methods for this annotation: KNN, SVM, scANVI, and scGPT. For cells with known cell types, we divided the data into a training set (90%) and a holdout set (10%). The final cell type annotations were determined using a majority-vote system, where a cell type was confirmed only if at least two of the four methods agreed. Cells that did not meet this criterion were labeled as “unclear.”

Post-annotation, we examined the clusters for consistency in cell type determination. All clusters demonstrated over 95% agreement in cell-type identification, allowing us to assign a predominant cell type to each cluster confidently. The NVU Atlas was then constructed by subsetting the integrated dataset to include only the NVU cell types, neurons, astrocytes, endothelial cells, and pericytes. Given the high proportion of neuronal data within the NVU Atlas, we further annotated neuronal subtypes. For this, we used similar annotation methods as before, with the BICCN cell ontology from the SEA-AD dataset serving as the reference set.

### Analysis of differential expression and Cell-Cell interactions in NVU cell types

Differential expression (DE) analyses were conducted using negative binomial mixed model implemented in the NEBULA R package (version 1.5.4)(35) using NEBULA-LN method that can accurately estimate a very large subject-level overdispersion. We filtered low expressed genes where the counts per cell using threshold cpc=0.005 (i.e., counts per cell < 0.5%). A model was constructed as diagnosis + age at death + source. Genes failed to converge were removed by requiring convergency > −20. P-values were adjusted by Benjamini-Hochberg procedure for multiple comparison correction. Genes with |logFC| > 0.2 and adjusted p-value < 0.05 were selected as significant DE genes. Pathway analysis was performed with R package enrichR(40) against KEGG_2021_Human database.

Possible cell-cell interactions (CCI) between NVU cell types were identified within the DE gene set using a manually curated ligand-receptor (LR) database that integrates four databases: CellChat(36), CellPhoneDB(37), CellTalkDB(38), and SingleCellSignalR(39). LR pairs were selected if both ligand and receptor were differentially expressed in at least one cell type. These pairs were ranked by the product of their log fold changes.

### Database and its deployment

We have implemented a user-friendly interface with RShiny and hosted at: https://rtools.mayo.edu/NVU_Atlas/.

## RESULTS

### Comprehensive list of data sets used for the development of the Human NVU Atlas

The Human NVU Atlas is compiled from 11 Alzheimer’s-related studies from public databases, GEO, Synapse, and CellxGene. While most studies focus on general brain functions and Alzheimer’s-related changes, predominantly profiling neurons, three studies are specifically enriched for brain vasculatures, enhancing representation of rarer blood-brain barrier cell types like endothelial cells and pericytes in the NVU atlas. These 11 datasets include single-nuclei transcriptome data from approximately 4,252,206 cells derived from 748 patients, balanced in terms of sex and disease condition as detailed in **Table 1**. All studies we focused to develop the NVU atlas targeted the pre-frontal cortex and employed droplet-based 10X 3’ sequencing technology, ensuring a uniform approach to data generation. Additionally, the use of Chromium Single Cell 3’ v3 sequencing technology across all studies enhances the depth and reliability of the analyses.

### Heterogeneity and integration in cell type distribution of the Prefrontal Cortex

Despite the consistent focus on the pre-frontal cortex, we observed significant heterogeneity in cell type abundance across the integrated cohorts. This variance primarily is observed from the differing emphases of each study. **Figure 1a** displays the 11 publicly available sources of cell samples used in the pre-frontal Atlas, with each color representing a different resource/dataset, illustrating the extensive integration effort. **Figure 1b** details the cell type distribution within the atlas, categorizing cells into major types such as neurons, astrocytes, endothelial cells, and others, including about 25% of cells classified as unknown types. The color coding in **Figure 1b** highlights the atlas’s diverse cell types and the curated assembly, with nearly 75% of data featuring author-provided cell-level annotations, offers valuable insights into the cellular and molecular architecture of the frontal cortex in Alzheimer’s Disease. However, to annotate the unknown cells depicted in **Figure 1b**, we employed four methods: two traditional machine learning approaches, KNN and SVM, which have shown good performance in benchmark studies, and two deep learning-based methods, scANVI and scgpt. A 10-fold validation assessed each method’s effectiveness, with KNN, SVM, and scgpt demonstrating over 99% accuracy in both training and holdout datasets, while scANVI achieved 97% precision but a lower recall of 69%. We then applied all four models to the 25% of data with unknown cell types, and the final cell type classification was determined by majority agreement among the methods.

**Figure 1:**
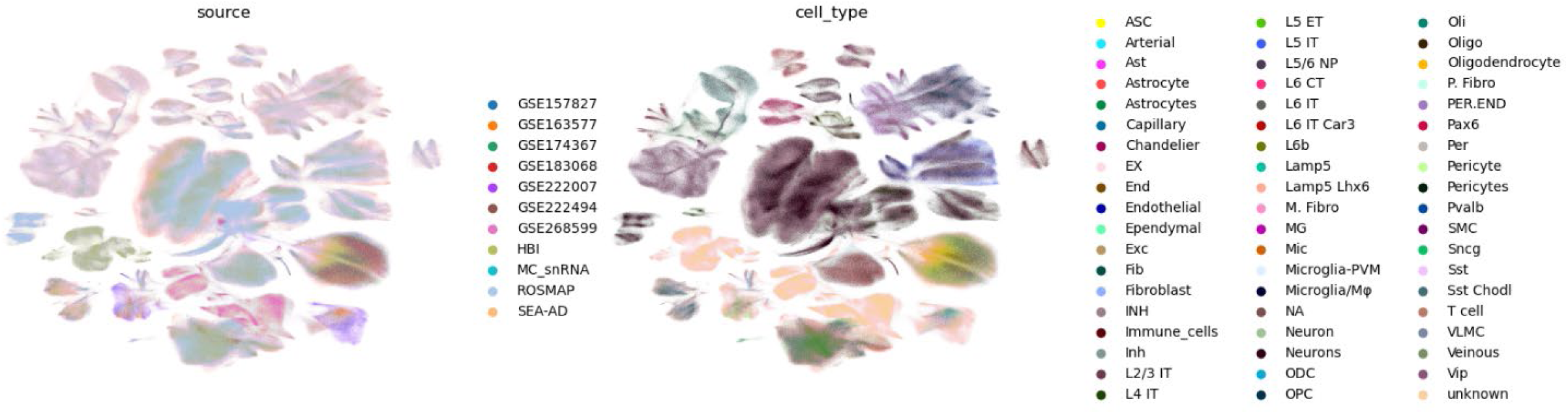
Visualization of cell sources and cell type distributions in the Human NVU Atlas. **(a)** This visualization shows the distribution of cell sources in the Human NVU Atlas. Each cluster represents cells derived from different studies as indicated by the color legend. The mixing of colors across clusters suggests the integration of data from multiple sources, highlighting the comprehensive nature of the atlas. (**b**) This image illustrates the major cell type distribution within the Human NVU Atlas. Distinct colors represent different cell types as detailed in the color legend, showing the diversity and density of cell types such as neurons, astrocytes, endothelial cells, and others. This visual representation emphasizes the utility of the atlas in providing detailed insights into cell type-specific contexts within the neurovascular unit and broader neurological studies.

### Composition and cell type diversity in the integrated Human NVU Atlas

As shown in **Figure 2a**, following the integration and subsequent cell type annotation, the integrated Human NVU Atlas now encompasses single-nuclei transcriptome data from 3,352,997 nuclei, specifically covering various NVU cell types, neurons, astrocytes, endothelial cells, and pericytes. The composition of the atlas predominantly consists of neurons, which make up 80.7% of the total, followed by astrocytes at 14.6%. Endothelial cells and pericytes are less abundant, representing 2.8% and 1.9% of the atlas, respectively, as detailed in **Figure 2b**. Figures **2c and 2d** display the UMAP visualization of integrated cell clustering of the cell types. We further explored neuron subtypes using BCCIN cell type ontology through transfer-labeling from the SEA-AD dataset, as shown in **Figure 2d**. The neurons were categorized into various groups: ExcNeuron1 (L2/3 IT), ExcNeuron2 (L4 IT), ExcNeuron3 (L5 IT, L6 IT, and L6 Car3), ExcNeuron4 (other L5/L6 excitatory neurons), and inhibitory neurons. Particularly, the grouping of excitatory neurons aligns well with the layer origins of the neurons, indicating a consistent and biologically relevant classification.

**Figure 2:**
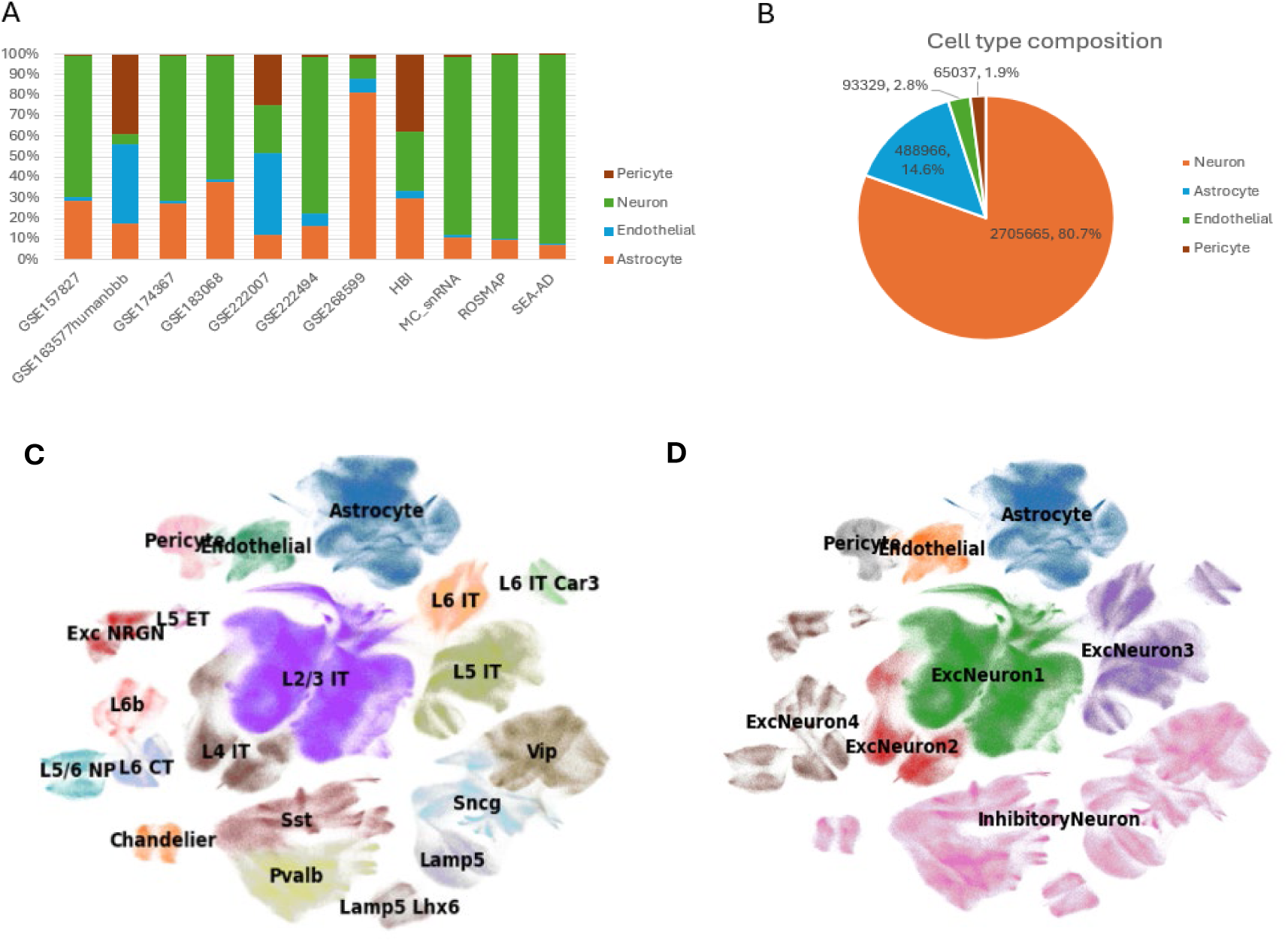
Integration and cell type annotation in the NVU Atlas. **(a)** Histogram displaying the proportion of cell types across different studies included in the Human NVU Atlas. Each bar represents a study and shows the percentage of pericytes, neurons, endothelial cells, and astrocytes. (**b**) Pie chart illustrates the overall cell type composition in the Human NVU Atlas. The chart highlights the dominance of neurons (80.7%), followed by astrocytes (14.6%), endothelial cells (2.8%), and pericytes (1.9%). (**c**) Uniform manifold approximation and projection (UMAP) visualization of integrated cell clustering within the Human NVU Atlas, annotated with major cell types including neurons, astrocytes, endothelial cells, and pericytes. (**d**) Detailed clustering of neuron subtypes within the Human NVU Atlas, identified by transfer-labeling BCCIN cell types from the SEA-AD dataset. This map categorizes neurons into ExcNeuron1 (L2/3 IT), ExcNeuron2 (L4 IT), ExcNeuron3 (L5 IT, L6 IT, and L6 Car3), ExcNeuron4 (other L5/L6 excitatory neurons), and inhibitory neurons, showing consistency with neuronal layer origins.

This atlas will facilitate understanding of the neurovascular unit’s role in Alzheimer’s pathology, highlighting specific cellular interactions and gene expression patterns that could potentially be targeted for therapeutic interventions. We have applied three use cases to demonstrate the utility of the Human NVU Atlas. We used the atlas to derive disease-specific pathways that are disrupted in individual NVU cell types, sex differences, and cell-cell interactions in NVU cell types and cell-type expression of genes in NVU cell types.

### Use Case #1: Cell-Type Specific Gene Expression Profiling in the Human NVU Atlas

The utilization of the Human NVU Atlas is illustrated through the analysis of cell-type specific gene expression within the neurovascular unit. By examining the expression patterns of specific genes, we can gain insights into their roles in different cell types associated with the NVU. In **Figure 3**, we showcase the expression patterns of four key genes: PECAM1, GFAP, PDGFRB, and RBFOX3 across different NVU cell types, using UMAP plots. These plots illustrate the specific molecular signatures of different NVU cell types, essential for understanding their roles in brain health and disease.

**Figure 3:**
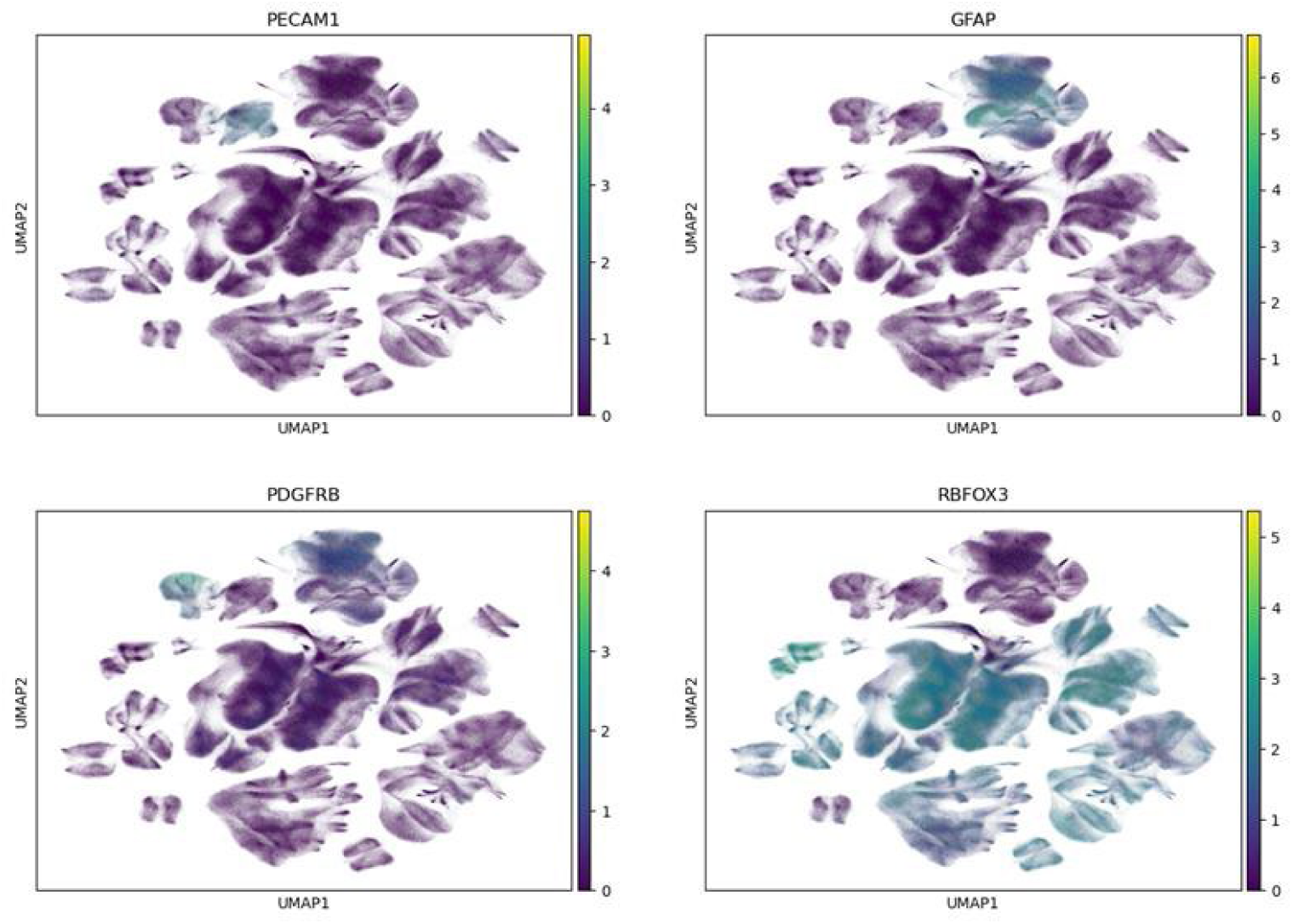
Cell-type specific gene expression in the Human NVU Atlas. UMAP plots depicting the expression of key genes across NVU cell types using the Human NVU Atlas. Each plot is color-coded to represent gene expression levels: **PECAM1**: Predominantly expressed in endothelial cells. **GFAP**: Highly expressed in astrocytes. **PDGFRB**: High expression in pericytes. **RBFOX3 (NeuN)**:Extensively expressed in neurons.

### PECAM1 Expression

The UMAP plot for PECAM1 highlights its expression predominantly within endothelial cells. This gene is crucial for endothelial cell function and is known to play a significant role in the vascular components of the NVU.

### GFAP Expression

This gene is a marker for astrocytes and is expressed strongly in these cells, as shown in the corresponding UMAP plot. The distribution shows the astrocytes’ role in the NVU, emphasizing their involvement in both the support of neuronal function and the maintenance of the blood-brain barrier.

### PDGFRB Expression

The expression of PDGFRB is seen primarily in pericytes, which are critical for blood vessel stability and blood-brain barrier integrity within the NVU. The visualization indicates a concentrated expression in specific clusters, reflecting the specialized functions of these cells in vascular health and pathology.

### RBFOX3 Expression

Also known as NeuN, this gene is a neuronal marker. The plot shows high expression levels in neuronal populations, highlighting its utility in identifying and studying neuronal cells within complex tissue environments like the NVU.

In summary, Use Case #1 shows the cellular dynamics within the neurovascular unit in Alzheimer’s disease, as revealed through cell-type specific gene expression profiling using the Human NVU Atlas. By mapping key genes such as PECAM1, GFAP, PDGFRB, and RBFOX3 across different cell types, we have deepened our understanding of how various cells contribute uniquely to the pathophysiology of AD. This use case highlights the critical roles played by endothelial cells, astrocytes, pericytes, and neurons in maintaining brain integrity.

### Use Case #2: Differential Gene Expression Analysis of cell types in the NVU for Alzheimer’s Disease

The analysis of differential gene expression (DGE) across various cell types in the neurovascular unit is a critical aspect of understanding cellular responses in the context of Alzheimer’s disease. In this use case, we explored gene expression changes within the neurovascular unit in Alzheimer’s disease compared to controls, focusing on key NVU cell types: excitatory and inhibitory neurons, astrocytes, endothelial cells, and pericytes. Each cell type was analyzed for differential expression, adjusting for sex, diagnosis, age, APOE status and sample source. This analysis included an FDR < 0.05 and a log fold change (logFC) > 0.2 for DGE.

### Gene Expression Analysis by Cell Type

We observed distinctive DGE patterns across cell types. The number of DEGs varied considerably across cell types (p=0.005, ANOVA test), with endothelial and pericyte populations exhibited more upregulated genes, whereas excitatory and inhibitory neurons showing the largest numbers of downregulated genes (Figure 4a, Supplementary Table S1). We did not observe numbers of DE genes with statistical difference between the sexes. Volcano plots further illustrate these patterns, with clear clusters of significantly downregulated genes in excitatory neurons and enriched upregulated signals in endothelial and pericyte populations (Figure 4b).

**Figure 4:**
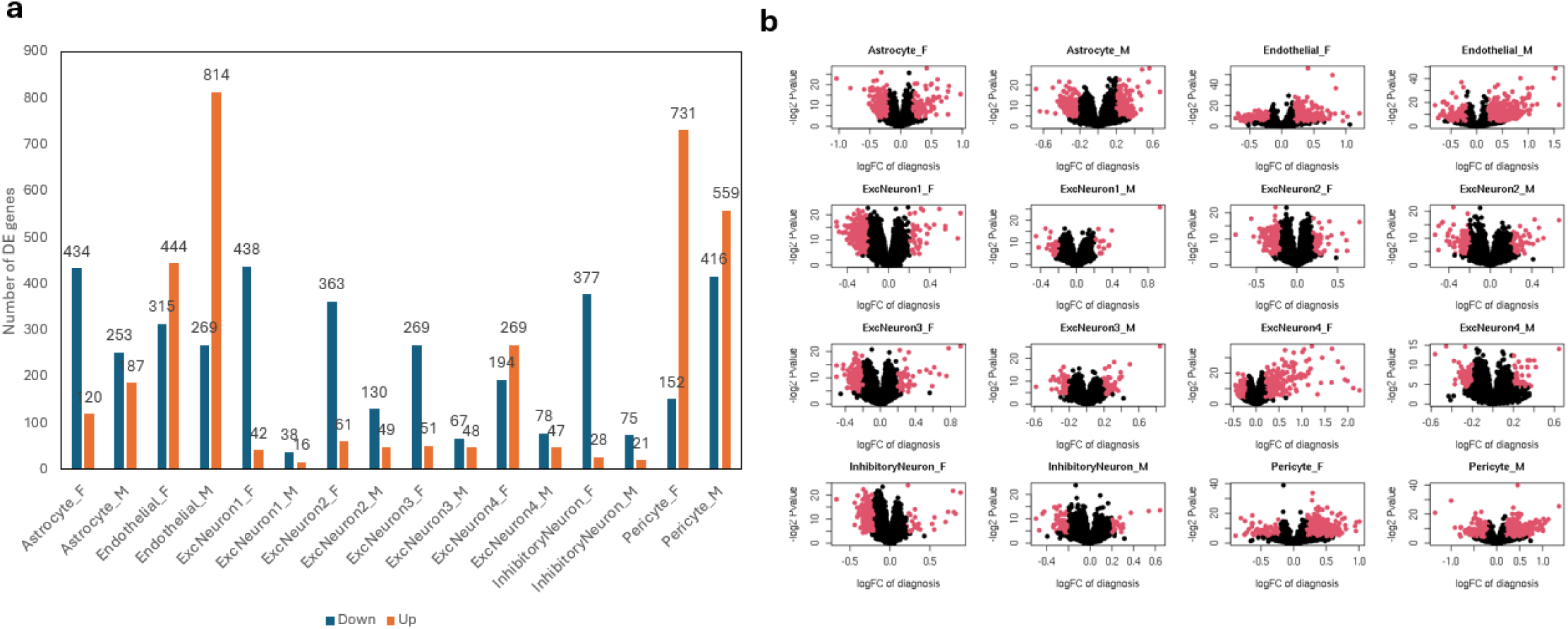
Differential Gene Expression Analysis in NVU Cell Types in Alzheimer’s Disease. (**a**) presents a summary table listing the total genes and differentially expressed genes in various NVU cell types by sex in Alzheimer’s disease versus control comparisons. (**b**) displays corresponding volcano plots for selected cell types, illustrating the logFC for the genes. Red points highlight genes that are significantly differentially expressed, with a focus on showing the variability and sex-specific expression patterns among excitatory neurons, inhibitory neurons, astrocytes, endothelial cells, and pericytes.

In astrocytes, we detected 434 downregulated and 120 upregulated genes in females, while male astrocytes showed 253 downregulated and 187 upregulated genes. We identified 121 common DE genes between male and female changing in the same direction, including elevated expression in CREB5 and NOTUM, both are potential therapeutic targets for AD (41,42). Commonly affected pathways including WNT signaling pathways, GABAergic synapse, MAPK signaling pathway, etc (Supplementary Table S2). In female, we observed enriched proinflammation pathways and down regulated gap junction pathway and insulin secretion. In male we observed down-regulated AGE-RAGE signaling pathway in diabetic complications.

Endothelial cells demonstrated pronounced transcriptional changes, with 1083 DE genes in male and 759 in female. The 140 common DE genes both sexes showed enrichment in upregulation of metallothioneins (MTs), insulin secretion related AD marker genes (SNAP25, ADCYAP1, and CCK). We also observed overlapped down-regulated pathways (Gap junctions and GABAergic synapse) in both male and female while through different genes, showing possible different mechanisms. In female we observed down regulation of cAMP signaling pathway, neuroactive ligand-receptor interaction, and insulin secretion. In male we observed up-regulation of insulin secretion and proinflammatory pathways, such as TNF signaling pathways.

Pericytes displayed highest number of DE genes across all cell types while it represented less than 2% of all cells in the whole atlas, with 975 in male and 883 in female. In the common DE genes, we observed enrichment in down regulation of insulin secretion genes and IL6. In the sex specific DE genes, we observed down-regulation of COL4A1 and COL4A2 only in female, which are known critical for BBB integrity and was not observed in the male pericytes. We also observed GAP junction up regulated in female but down regulated in male.

Common upregulated DE across the excitatory and inhibitory Neuron clusters suggested neurodegeneration and neuronal damage associated genes, such as upregulation of ITGA10, SLC25A18 and B2M. In common down regulated genes, we observed neuroprotective gene VGF, and neuronal differentiation-associated ETV5. Across the various neuronal cell clusters, we observed significant more DE genes in female than male (p=0.0014). Specifically, in female only DE genes, we observed down-regulation of Brain Expressed X-linked genes (BEX1, BEX2, BEX3, and BEX5) on chr X, cyclooxygenase genes (COX5B1, COX6B1, and COX7C) and Gap junction related genes (GNG3 and NDUFA1). We also observed female specific DE genes related to neuroinflammation (INHBA, CXCL13 and FOS).

### Pathway-Level Insights and Sex-Specific Differences

We extended our analysis to include pathway-level insights using the KEGG_human_2021 database through the R package enrichR, with findings detailed in Supplementary Table S2. Top 15 most frequently enriched pathways across the neurovascular unit was presented in Figure 5 for female and male, respectively.

**Figure 5.**
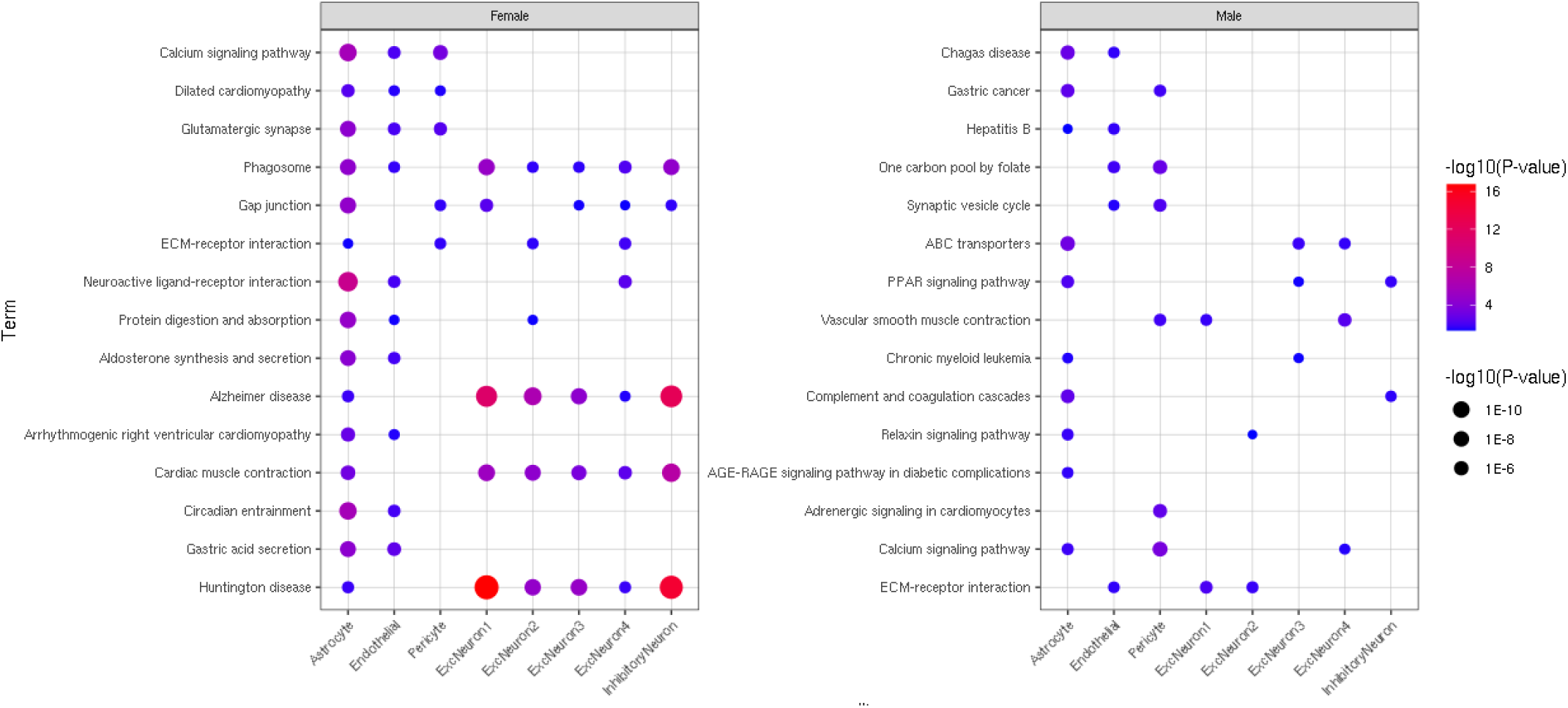
Top 15 most frequent pathways across cell types in female and male. The size and color of the dot indicates the significance in the pathway enrichment analysis, where red/large dots indicated higher significance.

We noticed that female showed more pathways being altered than male. Significant enrichment of ECM-receptor interactions and Calcium signaling pathway was observed in astrocyte, pericyte, and neurons across both sexes. We also noticed downregulation of insulin secretion in female astrocyte, endothelial and pericytes, which was also observed in endothelial and pericytes in male.

Astrocytes were the most altered pathways in both male and female. In female we observed enrichment in Gap junction, Alzheimer’s disease, neuroactive ligand-receptor interactions, while in male we noticed synaptic vesicle cycle and PPAR signaling pathway.

### Use Case #3: Sex-Specific Cell-Cell Communication Analysis in Alzheimer’s Disease

In our third use case, we focused on elucidating sex-specific differences in cell-cell communication within the NVU of Alzheimer’s disease patients compared to the controls. By analyzing potential ligand-receptor (LR) interactions, we aimed to uncover how intercellular communication varies between male and female patients and how these differences might influence disease progression.

**Figure 6** presents a comprehensive view of the identified potential LR interactions segregated by sex. It is noticed that endothelial cells were the most active signal initiators and receivers across AD brains. Pericytes and astrocytes were also major signaling hubs, engaging in extensive bidirectional communication with each other and with neurons (Figure 6a-b).

**Figure 6:**
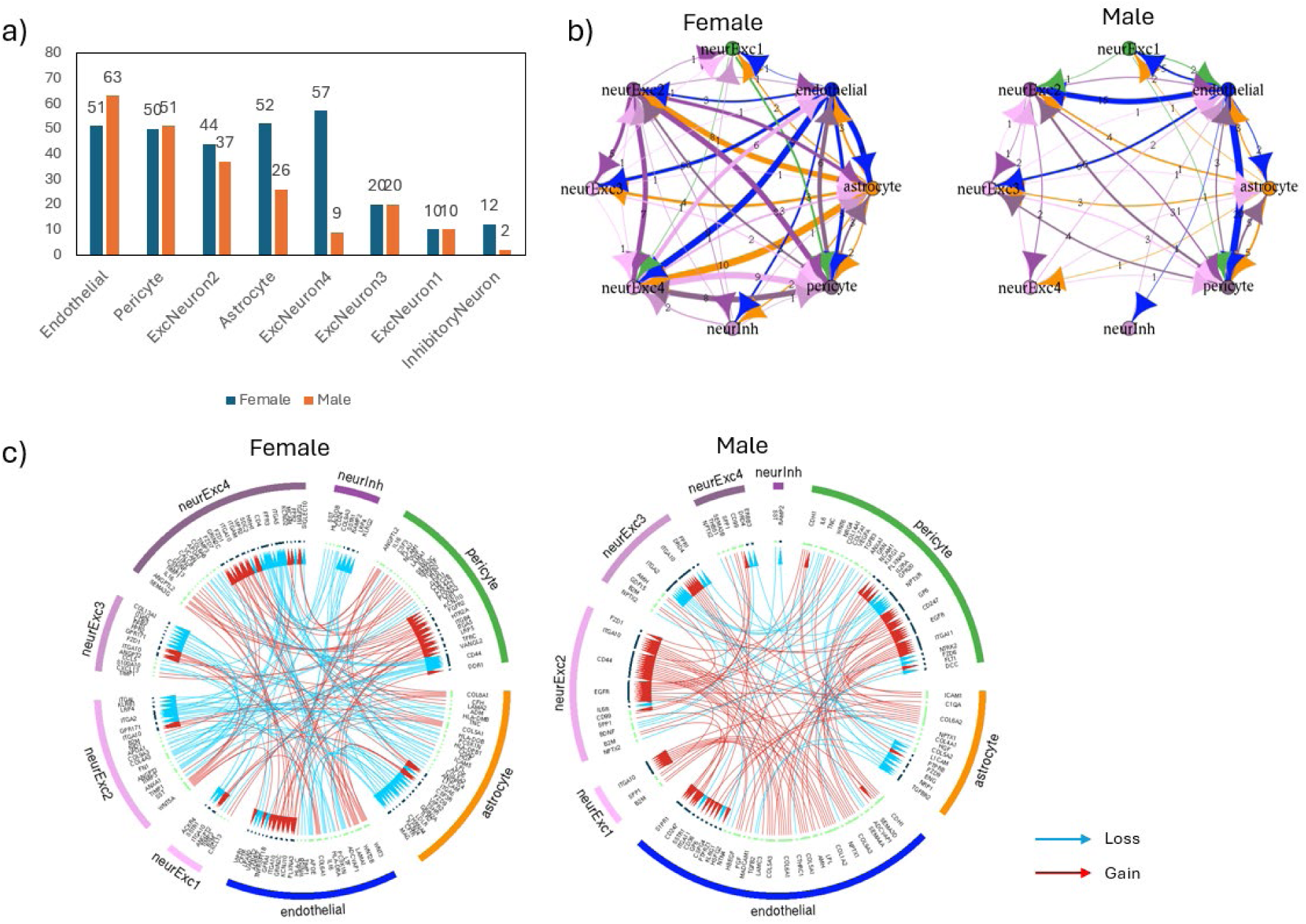
Sex-specific ligand-receptor interaction networks in Alzheimer’s disease compared to controls. **a.** The number of CCI candidates identified across cell types, sorting in descending order. **b.** A NetView diagram to show the interactions across cell types. **c**. Illustration of the sex-specific ligand-receptor (LR) interaction networks within the NVU of AD patients compared to controls, separated into female (left) and male (right) profiles. Each circular plot maps potential LR interactions across various NVU cell types including neurons, astrocytes, endothelial cells, and pericytes. Blue arrow arches represent loss of LR interaction and red represent gain, initiating from the ligands pointing to the receptors.

The circos LR plot showed key LR changes related to BBB integrity, inflammation, and metabolism in both sexes (Figure 6c). The analysis revealed common loss in collagen IV (COL4A1)-integrin alpha (ITGA2) between pericytes and excitatory neurons in both males and females (shown in blue). As a key collagen protein on the basement membrane, this breakdown could potentially compromise the integrity of the BBB and small blood vessels in the brain. Additional loss of BBB supportive LR interactions were observed in female. For example, another key base membrane LR pair COL4A2-ITGA2 showed as loss. Mixed signals of other collagens were observed in both sexes.

The changes in inflammation-related LR interactions (shown in red) were observed in both sexes, highlighting an overall dysregulation of inflammatory pathways in AD. However, the specific mediators of these interactions differed markedly between sexes. In female, we observed the gain of LR interactions of angiopoietin-like protein ANGPTL2 and integrin protein ITGA5 among pericytes, astrocytes and excitatory neurons, which possibly promoted neuroinflammation (43,44). While the LR interactions of angiopoietin ANGPT2 and integrin ITGA5 were observed to be in loss, which indicates loss of neuroprotective environment (45). Additional increased LR signaling across cell types include CXCL13-HTR2A, showing increased response to neuroinflammation. For male, reduced IL6–IL6R (pericytes to neurons) and TGFB2/3-TGFB2/ENG (from vasculature types to astrocytes) were observed, indicating cytokine suppression and reduced anti-inflammatory regulation.

Further, APOE-LRP4/LDLR interactions between vasculature types and neurons were found reduced only in females, showing possibly impaired clearance of amyloid-beta peptides.

## Discussion

In this study, we constructed the Human NVU Atlas, a comprehensive transcriptomic resource designed to resolve the cellular heterogeneity of the NVU in the human prefrontal cortex. By integrating data from 11 publicly available cohorts, including specific vascular-enriched datasets, we compiled over 4.25 million nuclei from 748 individuals. This integration addresses a critical gap in current AD research: the underrepresentation of vascular cells in standard snRNA-seq studies due to technical biases favoring neuronal recovery. By employing a unified probabilistic pipeline and advanced machine learning re-annotation strategies, we successfully recovered and identified substantial populations of endothelial cells (2.8%) and pericytes (1.9%), alongside the more abundant neuronal and astrocytic populations. Our NVU atlas is the largest resource consisting of 158,366 cells to study endothelial cell and pericyte interactions in Alzheimer’s disease.

Our differential gene expression analysis offers compelling transcriptomic evidence supporting the significant role of vascular dysfunction in AD pathogenesis. Despite their relatively low abundance compared to neurons, endothelial cells and pericytes exhibited a disproportionately high number of upregulated genes in AD patients. This suggests that while neurons may undergo degeneration and downregulation of key functional genes, the vascular components of the NVU enter a state of transcriptional hyperactivity, potentially reflecting a stress response or compensatory mechanism. This aligns with previous literature suggesting that BBB dysfunction is an early driver of neurodegeneration rather than merely a downstream consequence (9,10).

A defining insight from the NVU Atlas is the pronounced sexual dimorphism observed in AD pathology. Our analysis revealed that female brains exhibit significantly more altered pathways than males across the NVU. We observed distinct inflammatory signatures between sexes.

Females showed a gain in ligand-receptor interactions involving angiopoietin-like protein ANGPTL2 and integrin ITGA5, suggesting a pro-inflammatory environment. In contrast, males exhibited a reduction in IL6–IL6R signaling, indicating cytokine suppression. While both sexes showed a loss of collagen interactions (e.g., COL4A1-ITGA2) indicative of basement membrane breakdown, females uniquely displayed downregulation of specific collagen genes (COL4A1/2) in pericytes, which are critical for maintaining BBB integrity. We noted a downregulation of insulin secretion pathways in female astrocytes, endothelial cells, and pericytes, pointing toward metabolic dysregulation as a significant component of AD in women. These findings underscore the necessity of sex-stratified approaches in AD therapeutics, as the molecular drivers of disease progression appear distinct between males and females.

Our analysis also reveals the disrupted cell-cell communication in the NVU. The NVU functions as an integrated and interdependent system, and our interactome analysis highlights how this cohesion fractures in AD. We identified a widespread disruption in paracrine signaling, particularly involving the basement membrane, which acts as the structural anchor for the NVU. The consistent loss of collagen-integrin interactions between pericytes and excitatory neurons suggests a mechanical and functional uncoupling of the neurovascular unit. Furthermore, the reduction in APOE-LRP4/LDLR interactions observed specifically in females may indicate impaired clearance of amyloid-beta, providing a potential mechanistic link between the *APOE* genotype, sex, and vascular clearance efficiency.

### Limitations

While the NVU Atlas represents a significant advancement, it is not without limitations. First, the reliance on snRNA-seq means that cytoplasmic transcripts and local mRNA pools are not captured. Complex polarized cells from the cytoplasm might miss from the scene. Second, while we integrated vascular-enriched datasets, the inherent difficulty in isolating fragile vascular nuclei from post-mortem tissue means these cells remain rarer than in their native state. Finally, the current atlas focuses exclusively on the prefrontal cortex; future iterations will need to expand to other brain regions to capture the spatial heterogeneity of AD pathology.

## Conclusion and Future Directions

The Human NVU Atlas serves as a foundational resource for the AD research community, enabling systematic investigation of the non-neuronal contributions to cognitive decline. By providing high-confidence annotations and identifying novel sex-specific ligand-receptor disruptions, this work provides great resources for therapeutic targeting research of the BBB and neuroinflammation. We envision this atlas acting as a reference for deconvolution in spatial transcriptomics and bulk RNA-seq studies, furthering our ability to map the complex landscape of the Alzheimer’s brain. To facilitate convenient community access, a user-friendly interactive web-based interface is also provided at https://rtools.mayo.edu/NVU_Atlas/.

## Acknowledgement

This work was supported by National Institutes of Health/National Institute on Aging [R01AG085900] and National Institutes of Health/National Institute of Neurological Disorders and Stroke [R01NS125437].

